# Boceprevir, calpain inhibitors II and XII, and GC-376 have broad-spectrum antiviral activity against coronaviruses in cell culture

**DOI:** 10.1101/2020.10.30.362335

**Authors:** Yanmei Hu, Chunlong Ma, Tommy Szeto, Brett Hurst, Bart Tarbet, Jun Wang

## Abstract

As the COVID-19 pandemic continues to fold out, the morbidity and mortality are increasing daily. Effective treatment for SARS-CoV-2 is urgently needed. We recently discovered four SARS-CoV-2 main protease (M^pro^) inhibitors including boceprevir, calpain inhibitors II and XII and GC-376 with potent antiviral activity against infectious SARS-CoV-2 in cell culture. Despite the weaker enzymatic inhibition of calpain inhibitors II and XII against M^pro^ compared to GC-376, calpain inhibitors II and XII had more potent cellular antiviral activity. This observation promoted us to hypothesize that the cellular antiviral activity of calpain inhibitors II and XII might also involve the inhibition of cathepsin L in addition to M^pro^. To test this hypothesis, we tested calpain inhibitors II and XII in the SARS-CoV-2 pseudovirus neutralization assay in Vero E6 cells and found that both compounds significantly decreased pseudoviral particle entry into cells, indicating their role in inhibiting cathepsin L. The involvement of cathepsin L was further confirmed in the drug time-of-addition experiment. In addition, we found that these four compounds not only inhibit SARS-CoV-2, but also SARS-CoV, MERS-CoV, as well as human coronaviruses (CoVs) 229E, OC43, and NL63. The mechanism of action is through targeting the viral M^pro^, which was supported by the thermal shift binding assay and enzymatic FRET assay. We further showed that these four compounds have additive antiviral effect when combined with remdesivir. Altogether, these results suggest that boceprevir, calpain inhibitors II and XII, and GC-376 are not only promising antiviral drug candidates against existing human coronaviruses, but also might work against future emerging CoVs.

---

Coronaviruses (CoVs) are enveloped positive-stranded RNA virus which infect humans and multiple species of animals, causing a variety of highly prevalent and severe diseases.^*1, 2*^ In the last two decades, three highly pathogenic and lethal human coronaviruses have emerged: severe acute respiratory syndrome coronavirus (SARS-CoV), the virus that caused the outbreak of severe acute respiratory syndrome in human in Southern China in 2002 and killed 774 people among 8,098 infected worldwide;^*3*^ MERS-CoV, which causes severe respiratory disease outbreak in Middle East in 2012, led to 791 deaths among 2,229 infected;^*4*^ the SARS-CoV-2, a novel coronavirus emerged in China in December 2019, quickly spread worldwide and became a global pandemic.^*5*^ As of October 28^th^ 2020, there are more than 44 million confirmed cases and over 1.1 million deaths worldwide and these numbers are increasing daily (https://coronavirus.jhu.edu/map.html). In addition, human coronavirus (HCoV) strains 229E (HCoV-229E), NL63 (HCoV-NL63), OC43 (HCoV-OC43), and HKU1 (HCoV-HKU1) cause a significant portion of annual upper and lower respiratory tract infections in humans, including common colds, bronchiolitis, and pneumonia.^*6–8*^ There are currently no antivirals or vaccines for either the highly pathogenic SARS-CoV-2, SARS-CoV, MERS-CoV or the human CoVs. The current COVID-19 pandemic is a timely reminder of the urgent need for therapeutics against coronavirus infection. As future coronavirus outbreak cannot be excluded, it is desired to develop broad-spectrum antivirals that are not only active against existing CoVs, but also future emerging CoVs.

Coronavirus genome ranges from 26 to 32□kb, of which the 3′-terminal region, proximately one-third of the genome, encodes a number of structural proteins (spike protein, envelope protein, membrane protein, and nucleocapsid protein), while the 5′-terminal region, approximately two-thirds of the genome, encodes for the non-structural proteins (3-chymotrypsin-like protease (3CL or main protease), papain-like protease, helicase, RNA-dependent RNA polymerase, exoribonuclease and endoribonuclease, methyl transferase), and accessary proteins.^*9*^ Among the viral proteins under investigation as antiviral drug targets, the main protease (M^pro^) appears to be a high-profile drug target for development of broad-spectrum antivirals for the following reasons: 1) The M^pro^ plays an essential role in coronavirus replication by cleaving the viral polyproteins at more than 11 sites;^*10*^ 2) The M^pro^s have relatively high sequence similarity within each CoV group;^*11*^ 3) The M^pro^ has an unique substrate preference for glutamine at the P1 site (Leu-Gln↓(Ser,Ala,Gly)), a feature that is absent in closely related host proteases,^*12*^ suggesting it is feasible to design M^pro^ inhibitors with a high selectivity; (4) The structures of M^pro^s from multiple members of CoV family have been solved,^*13–16*^ paving the way for rational drug design.

We recently discovered four SARS-CoV-2 M^pro^ inhibitors including boceprevir, calpain inhibitors II and XII, and GC-376 (Figure 1).^*17*^ They had single-digit micromolar to submicromolar IC_50_ values in the enzymatic assay, and inhibited SARS-CoV-2 viral replication in cell culture with EC_50_ values in the single-digit micromolar to submicromolar range.^*17*^ The co-crystal structures of GC-376, calpain inhibitors II and XII in complex with SARS-CoV-2 M^pro^ have been solved, providing a molecular explanation for the tight binding of these compounds towards M^pro^.^*17, 18*^ Interestingly, despite the weaker enzymatic inhibition potency of calpain inhibitors II and XII against M^pro^ (IC_50_ = 0.97 µM and 0.45 µM, respectively) compared with GC-376 (IC_50_ = 0.03 µM), calpain inhibitors II and XII had more potent cellular antiviral activity (EC_50_ = 2.07 µM and 0.49 µM, respectively) than GC-376 (EC_50_ = 3.37 µM). We also found calpain inhibitors II and XII inhibit human cathepsin L in the *in vitro* enzymatic assay,^*18*^ and cathepsin L has been shown to play an essential role in SARS-CoV-2 cell entry by activating the viral spike protein in the late endosome or lysosome.^*19, 20*^ These observations led us to speculate that the potent cellular antiviral activity of calpain inhibitors II and XII might be a result of inhibiting both viral M^pro^ and host cathepsin L. To test the hypothesis that the cellular antiviral activity of calpain inhibitors II and XII involves the inhibition of cathepsin L, we performed SARS-CoV-2 pseudovirus neutralization assay and the drug time-of-addition assay using HCoV-OC43 virus. Results from both assays indeed support the contribution of inhibition of cathepsin L to the cellular antiviral activity of calpain inhibitors II and XII.

**Figure 1.**
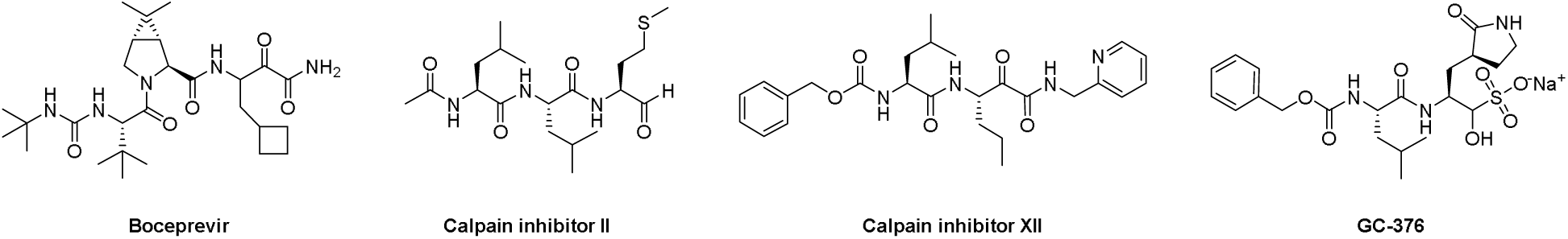
Chemical structures of SARS-CoV-2 M^pro^ inhibitors boceprevir, calpain inhibitors II and XII and GC-376.

To evaluate the broad-spectrum antiviral activity of boceprevir, calpain inhibitors II and XII, and GC-376, we tested them against two other highly pathogenic coronaviruses the SARS-CoV and MERS-CoV, as well as three human coronaviruses HCoV-OC43, HCoV-NL63, and HCoV-229E. The mechanism of action was studied by the thermal shift binding assay and FRET-based enzymatic assay. The combination therapy potential of these four compounds with remdesivir was also quantified by the drug combination index method. Altogether, our work demonstrated that boceprevir, calpain inhibitors II and XII, and GC-376 are promising drug candidates for the design and development of broad-spectrum antivirals against current and future emerging CoVs.

## RESULTS AND DISCUSSION

### Calpain inhibitors II and XII, but not GC-376 or boceprevir, blocked the entry of SARS-CoV-2 pseudoviral particles

Pseudovirus neutralization assay is an established model to study the mechanism of viral cell entry and has been widely used to assess the antiviral activity of viral entry/fusion inhibitors.^*21, 22*^ Calpain inhibitors II and XII are potent inhibitors of human cathepsin L, with K_I_ and IC_50_ values in the nanomolar range.^*18, 23*^ Cathepsin L plays an important role in SARS-CoV-2 viral entry by cleaving the viral spike S protein in the endosome or lysosome.^*19, 20*^ We hypothesized that cellular antiviral mechanism of calpain inhibitors II and XII might involve inhibition of cathepsin L besides M^pro^. To test this hypothesis, we first amplified SARS-CoV-2 pseudoviral particles in ACE2-expressing HEK293T cells (ACE2/293T) as previously described.^*21*^ We then performed pseudovirus entry assay in Vero E6 cells, which have been shown to have minimal levels of TMPRSS2 expression, therefore SARS-CoV-2 cell entry is mainly mediated through endocytosis, which relies on cathepsin L for viral spike protein activation.^*19*^ E-64d, a known cathepsin L inhibitor, was included as a positive control and it was found to inhibit SARS-CoV-2 pseudovirus entry with an IC_50_ value of 0.91 ± 0.10 µM (Figure 2A). Both calpain inhibitors II and XII also showed inhibitory activity against SARS-CoV-2 pseudovirus entry into Vero E6 cells with IC_50_ values of 9.26 ± 1.35 and 5.28 ± 0.74 µM, respectively (Figures 2B and 2C). However, neither GC-376 nor boceprevir had significant effect on SARS-CoV-2 pseudovirus entry at up to 100 µM concentration (Figures 2D and 2E). Altogether, it can be concluded that the cellular antiviral activity of calpain inhibitors II and XII involves the inhibition of cathepsin L, while GC-376 and boceprevir had no effect on cathepsin L-mediated viral entry.

**Figure 2.**
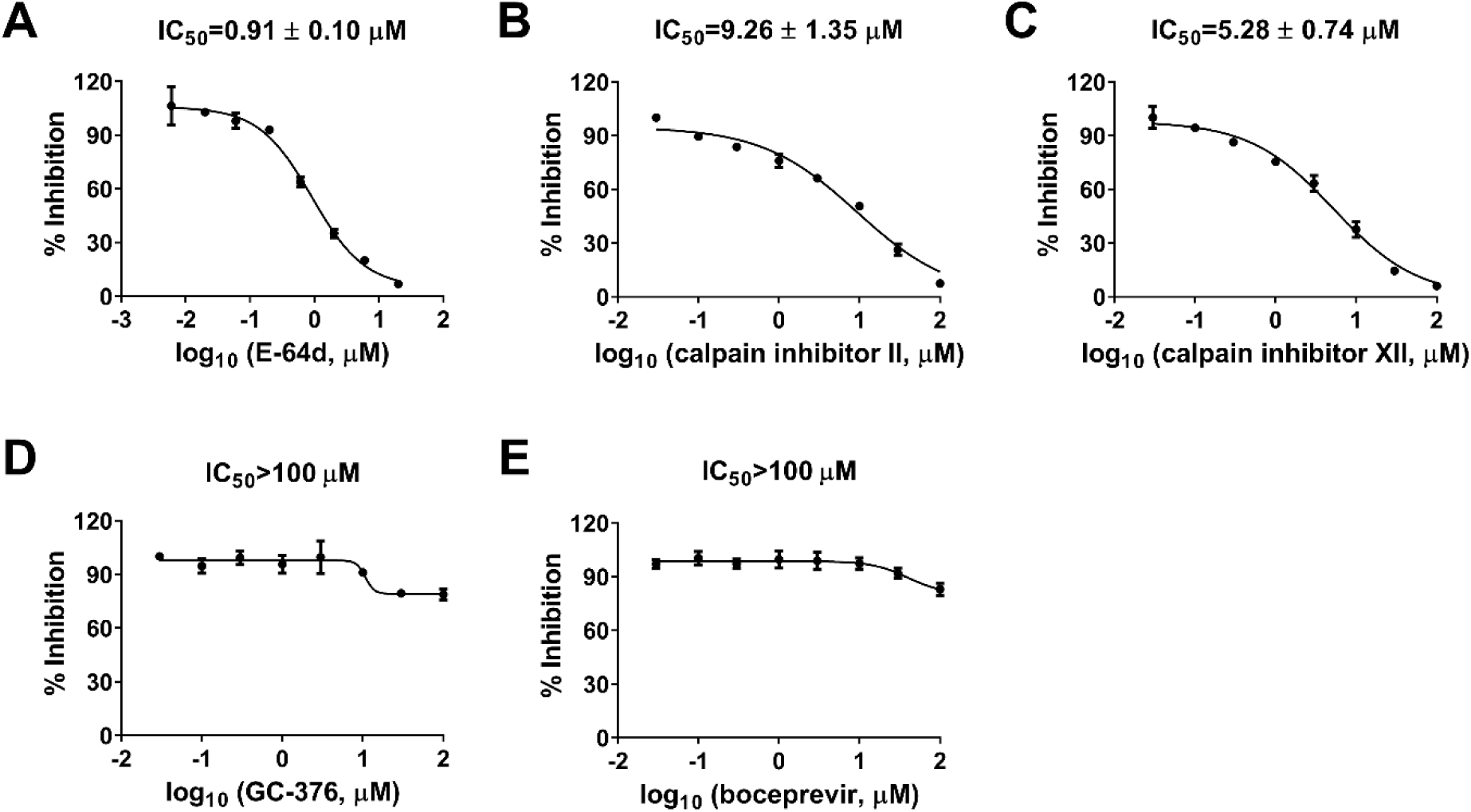
Inhibitory activity of boceprevir, calpain inhibitors II and XII, and GC-376 in the SARS-CoV-2 pseudovirus neutralization assay. Effect of E-64d (A), calpain inhibitor II (B), calpain inhibitor XII (C), GC-376 (D), and boceprevir (E) on SARS-CoV-2 viral entry in the pseudovirus assay. EC_50_ curve fittings using log (concentration of inhibitors) vs percentage of inhibition with variable slopes were performed in Prism 8. Data are mean ± standard deviation of two replicates.

### Drug time-of-addition experiment showed that calpain inhibitor XII inhibited HCoV-OC43 replication at both the early and intermediate stages of viral replication

M^pro^ mediates the cleavage of viral polyproteins pp1a and pp1ab during the viral replication process, therefore M^pro^ inhibitors such as GC-376 are expected to inhibit viral replication in the intermediate stage of viral replication in the drug time-of-addition experiment. In contrast, cathepsin L plays a role in the early stage of viral fusion by cleaving the viral spike protein. As such, cathepsin L inhibitor should inhibit the early stage of viral replication. Dual inhibitors such as calpain inhibitors II and XII are expected to inhibit viral replication at both the early and the intermediate stages. To test this hypothesis, we chose calpain inhibitor XII as a representative example and performed the drug time-of-addition experiment by adding inhibitor at different time points either before, during, or after HCoV-OC43 viral replication (Figure 3). GC-376 was included as a control of monospecific M^pro^ inhibitor. Viruses in the supernatant were harvested 16 hpi and the viral titers were quantified by plaque reduction assay. HCoV-OC43 was used for this experiment as it enters cells via endocytosis and relies on cathepsin L in late endosome for viral spike protein cleavage and activation,^*24*^ similar to SARS-CoV-2. HCoV-OC43 is a BSL-2 pathogen and is frequently used as surrogate virus of SARS-CoV-2 for mechanistic studies. Both viruses belong to the β-coronavirus lineage and share many similarities. When applied during viral infection and removed through washing afterward, GC-376 showed no antiviral activity (Figure 3), suggesting that GC-376 had no effect on virus entry. The antiviral potency of GC-376 gradually decreased when it was added at later stages of viral replication (6 and 9 hpi) (Figure 3), indicating that GC-376 inhibited viral replication at an intermediate stage, which is in line with its mechanism of action by targeting M^pro^. In contrast, calpain inhibitor XII inhibited both the early and intermediate stages of viral replication. Specifically, compared to DMSO control, ∼ 1 log_10_ decrease in the viral tier was observed when it was added during virus infection, suggesting it inhibited the early stage of viral entry (Figure 3). In addition, the antiviral potency of calpain inhibitor XII gradually decreased when added at later stages of viral replication (6 and 9 hpi) (Figure 3). Taken together, the drug time-of-addition experiment further confirmed the dual cellular antiviral mechanisms of action of calpain inhibitor XII by targeting both host cathepsin L and viral M^pro^.

**Figure 3.**
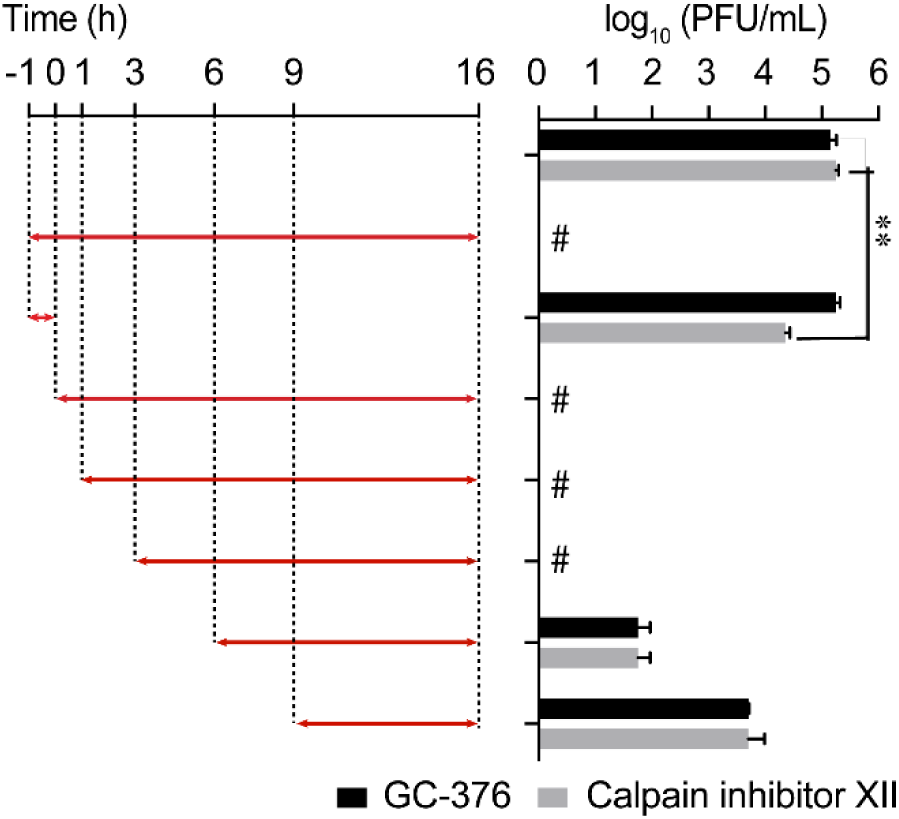
Time-of-addition profiles of GC-376 and calpain inhibitor XII. RD cells were infected with HCoV-OC43 virus at −1 h time point, viruses were incubated at 37 °C for 1 h for viral attachment and entry. At time point 0 h, cells were washed with PBS buffer twice and viruses from cell culture medium were harvested at 16 h p.i. The titer of harvested virus was determined by plaque assay. Arrows indicate the period in which 10 μM GC-376 or 25 μM calpain inhibitor XII was present. # indicates no virus was detected. Asterisks indicate statistically significant difference in comparison with the DMSO control (student’s t-test, **P < 0.005). The value of viral titer was the mean of two independent experiments ± standard deviation.

### Boceprevir, calpain inhibitors II and XII, and GC-376 directly bind to MERS-CoV, SARS-CoV and HCoV-OC43 M^pro^

First, we performed sequence alignment of M^pro^ from multiple members of coronavirus family: HCoV-229E, HCoV-OC43, HCoV-NL63, SARS-CoV, MERS-CoV, and SARS-CoV-2 (Figure S1). Overall, the M^pro^s showed moderate to high similarity in primary sequence, and comparatively high sequence similarity within each CoV group (Figure S1). It is acknowledged that 3-D structures of the M^pro^s are more conserved, especially at the active site.^*25*^ Therefore, we hypothesized that boceprevir, calpain inhibitors II and XII, and GC-376 might similarly bind to other CoV M^pro^s in addition to SARS-CoV-2 M^pro^. To test this hypothesis, we carried out differential scanning fluorimetry (DSF) assay.^*26*^ Specific binding of a ligand to a protein typically stabilizes the target protein, resulting in an increased melting temperature (*T*_m_). It was found that boceprevir, calpain inhibitors II and XII, and GC-376 increased the *T*_m_ of SARS-CoV, MERS-CoV and HCoV-OC43 M^pro^ in a dose dependent manner (Figure 4), while remdesivir had no effect on HCoV-OC43 M^pro^ stability at up to 200 µM. This was expected as remdesivir is a known RNA-dependent RNA polymerase (RdRp) inhibitor.^*27*^ GC-376 significantly increased the stability of all three M^pro^s when tested at 6 µM, with Δ*T*_*m*_ of 17.23, 9.78, and 13.86 °C against MERS-CoV, SARS-CoV and HCoV-OC43 M^pro^, respectively (Figure 4, Table 1). Boceprevir, calpain inhibitors II and XII also stabilized all three M^pro^s, but were less potent than GC-376. Boceprevir increased the *T*_*m*_ of MERS-CoV, SARS-CoV and HCoV-OC43 M^pro^ by 2.46, 3.94, and 1.02 °C, respectively at 60 µM (Figure 4, Table 1). Calpain inhibitors II and XII increased the *T*_*m*_ of MERS-CoV, SARS-CoV and HCoV-OC43 M^pro^ by 2.53, 2.88, 3.48 °C and 0.81, 2.41, 1.54 °C, respectively at 20 µM (Figure 4, Table 1). This result confirmed that boceprevir, calpain inhibitors II and XII, and GC-376 had direct binding towards SARS-CoV, MERS-CoV, and HCoV-OC43 M^pro^s in addition to SARS-CoV-2 M^pro^, indicating they might be broad-spectrum antiviral candidates.

**Figure 4.**
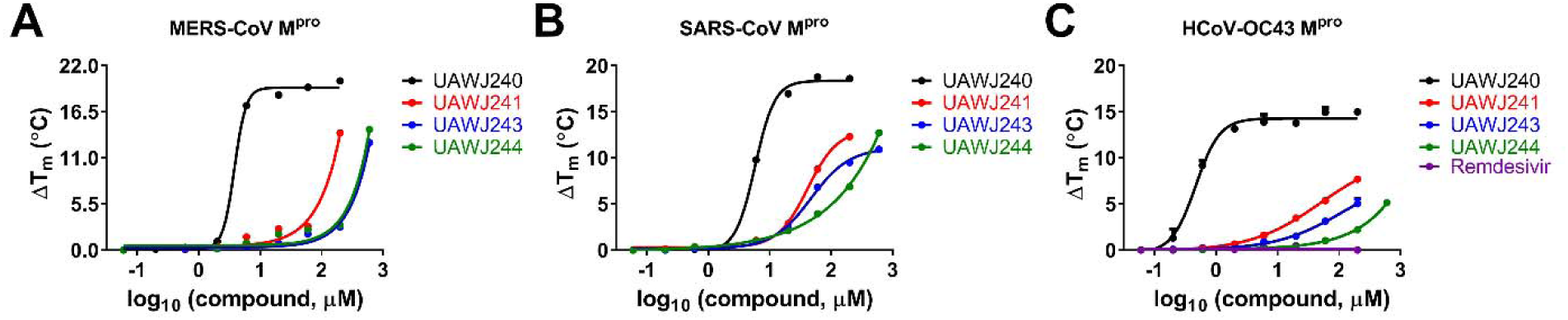
Effect of boceprevir, calpain inhibitors II and XII, and GC-376 on melting temperature (*T*_*m*_) of MERS-CoV M^pro^ (A), SARS-CoV M^pro^ (B), and HCoV-OC43 M^pro^ (C). Data were plotted with Δ*T*_*m*_ vs log_10_ (concentrations of compound) using Boltzmann Sigmoidal equation in Prism 8. Data are mean ± standard deviation of two replicates.

**Table 1.** Melting temperature shift (Δ*T*_m_) of MERS-CoV, SARS-CoV, and HCoV-OC43 M^pro^ in the presence of indicated concentrations of boceprevir, calpain inhibitors II and XII, and GC-376.

| Compounds | MERS-CoV $M^{pro}$<br>($\Delta T_m$, °C) | SARS-CoV $M^{pro}$<br>($\Delta T_m$, °C) | HCoV-OC43 $M^{pro}$<br>($\Delta T_m$, °C) |
| --- | --- | --- | --- |
| 200 μM remdesivir | N.T.<sup>a</sup> | N.T. | 0.00 ± 0.02 |
| 6 μM GC-376 | 17.23 ± 0.00 | 9.78 ± 0.07 | 13.86 ± 0.81 |
| 20 μM calpain inhibitors II | 2.53 ± 0.28 | 2.88 ± 0.21 | 3.48 ± 0.02 |
| 20 μM calpain inhibitors XII | 0.81 ± 0.14 | 2.41 ± 0.00 | 1.54 ± 0.01 |
| 60 μM boceprevir | 2.46 ± 0.07 | 3.94 ± 0.13 | 1.02 ± 0.05 |

<sup>a</sup>N.T. = not tested.

### Boceprevir, calpain inhibitors II and XII, and GC-376 inhibit the enzymatic activity of MERS-CoV, SARS-CoV and HCoV-OC43 M^pro^

To test whether boceprevir, calpain inhibitors II and XII, and GC-376 inhibit the enzymatic activity of other CoV M^pro^s, we performed enzyme kinetic studies of MERS-CoV, SARS-CoV and HCoV-OC43 M^pro^ with different concentrations of these four compounds (Figure 5). The enzymatic reactions were monitored for up to 4 h, and the progression curves are shown in Figures S2. As expected, similar to SARS-CoV-2 M^pro^, all four compounds showed biphasic progression curves at high drug concentrations, which are typical for slow covalent binding inhibitors. The first 90 min of the progression curves shown in Figure S2 were chosen for curve fitting as significant substrate depletion was observed when the proteolytic reaction proceeded beyond 90 min. The progression curves of all three CoV M^pro^ in the presence of different concentrations of GC-376 and MERS-CoV M^pro^ in the presence of different concentrations of boceprevir were fitted using the two-step reaction mechanism as previously described (Figure 5).^*17*^ In the first step, GC-376 binds to MERS-CoV, SARS-CoV and HCoV-OC43 M^pro^ with an equilibrium dissociation constant (K_I_) of 17.89 ± 2.34, 16.80 ± 2.36 and 3.63 ±0.26 nM, respectively (Table 2). After initial binding, a covalent bond is formed at a slower velocity in the second step between GC-376 and the M^pro^s with the second rate constant (k_2_) being 1.48, 0.87, and 0.31 s^−1^, respectively, resulting in an overall k_2_/K_I_ value of 82,910, 51,500 and 87,300 M^−1^ s^−1^, respectively (Figure 5A and Table 2). Boceprevir binds to MERS-CoV M^pro^ with a K_I_ of 1.65 ± 0.12 µM in the first step, and a k_2_ of 448.8 s^−1^ in the second step, resulting in an overall k_2_/K_I_ value of 272 M^−1^s^−1^ (Figure 5D and Table 2). When the proteolytic progression curves were fitted using the same two-step reaction mechanism, accurate values for the second rate constant k_2_ could not be obtained for calpain inhibitors II and XII against all three M^pro^s as well as boceprevir against SARS-CoV and HCoV-OC43 M^pro^s. This is presumably due to significant substrate depletion before reaching the equilibrium between EI and EI*, leading to very small values of k_2_. Therefore, only the dissociation constant K_I_ values from the first step were determined (Figure 5). The inhibition constants (K_I_) for calpain inhibitors II and XII with respect to MERS-CoV, SARS-CoV and HCoV-OC43 M^pro^ were 0.13 ± 0.012, 0.60 ±0.041, 0.23 ± 0.0088 µM and 1.32 ± 0.070, 0.14 ± 0.012, 0.43 ± 0.015 µM, respectively; while the K_I_ values for boceprevir with respect to SARS-CoV and HCoV-OC43 M^pro^ were 1.43 ± 0.14, and 2.29 ± 0.19 µM, respectively. Taken together, the enzymatic kinetic results suggest that boceprevir, calpain inhibitors II and XII, and GC-376 have broad-spectrum enzymatic inhibition against SARS-CoV, MERS-CoV, and HCoV-OC43 M^pro^.

**Figure 5.**
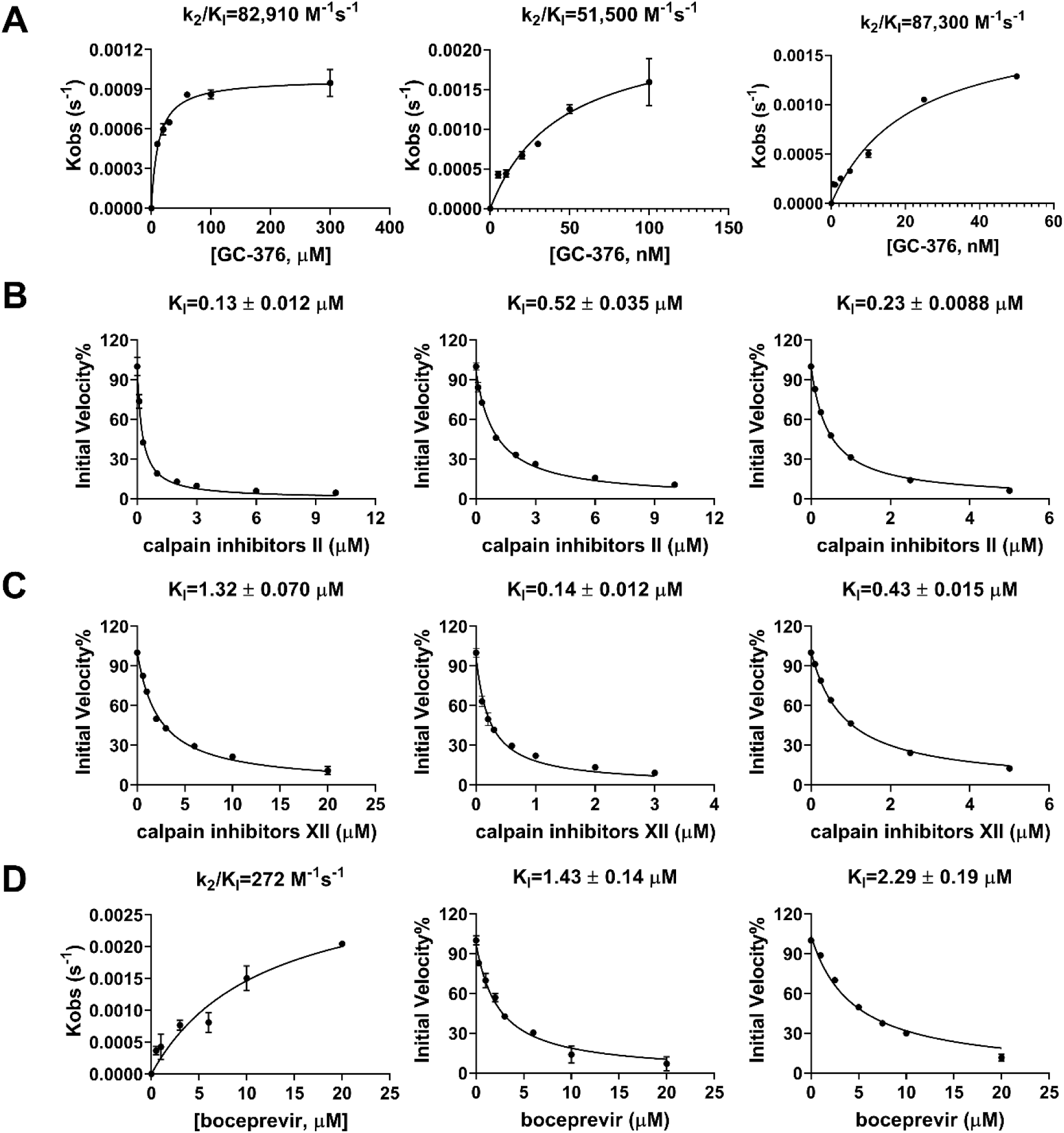
Data fittings of the proteolytic reaction progression curves of MERS-CoV M^pro^ (left column), SARS-CoV M^pro^ (middle column) and HCoV-OC43 M^pro^ (right column) in the presence or the absence of GC-376 (A); calpain inhibitor II (B); calpain inhibitor III (C); and boceprevir (D). In the kinetic studies, 60 nM MERS-CoV M^pro^, or 5 nM SARS-CoV M^pro^ or 3.3 nM HCoV-OC43 M^pro^ was added to a solution containing various concentrations of compounds and 20 µM FRET substrate to initiate the reaction. Detailed methods were described in “Materials and methods” section. Data are mean ± standard deviation of two replicates.

**Table 2.** Enzymatic inhibition of boceprevir, Calpain inhibitor II and XII, and GC-376 against various CoV M^pro^s.

| Compounds | SARS-CoV-2<sup>a</sup> | SARS-CoV | MERS-CoV | HCoV-OC43 |
| --- | --- | --- | --- | --- |
| GC-376 | IC<sub>50</sub>=0.030 ± 0.008 μM<br>k<sub>2</sub>/K<sub>I</sub> = 40,800 M<sup>-1</sup>s<sup>-1</sup> | IC<sub>50</sub>=0.079 ± 0.006 μM<br>k<sub>2</sub>/K<sub>I</sub> = 51,530 M<sup>-1</sup>s<sup>-1</sup> | IC<sub>50</sub>=0.110 ± 0.023 μM<br>k<sub>2</sub>/K<sub>I</sub> = 82,910 M<sup>-1</sup>s<sup>-1</sup> | IC<sub>50</sub>=0.011 ± 0.0012 μM<br>k<sub>2</sub>/K<sub>I</sub> = 87,300 M<sup>-1</sup>s<sup>-1</sup> |
| Calpain inhibitor II | IC<sub>50</sub>=0.97 ± 0.27 μM<br>K<sub>I</sub> = 0.40 μM | IC<sub>50</sub>=2.04 ± 0.20 μM<br>K<sub>I</sub> = 0.60 μM | IC<sub>50</sub>=1.24 ± 0.35 μM<br>K<sub>I</sub> = 0.13 μM | IC<sub>50</sub>=0.56 ± 0.08 μM<br>K<sub>I</sub> = 0.23 μM |
| Calpain Inhibitor XII | IC<sub>50</sub>=0.45 ± 0.06 μM<br>K<sub>I</sub> = 0.13 μM | IC<sub>50</sub>=0.60 ± 0.16 μM<br>K<sub>I</sub> = 0.14 μM | IC<sub>50</sub>=6.68 ± 2.76 μM<br>K<sub>I</sub> = 1.32 μM | IC<sub>50</sub>=0.34 ± 0.05 μM<br>K<sub>I</sub> = 0.43 μM |
| Boceprevir | IC<sub>50</sub>=4.13 ± 0.61 μM<br>K<sub>I</sub> = 1.18 μM | IC<sub>50</sub>=6.63 ± 1.05 μM<br>K<sub>I</sub> = 1.43 μM | IC<sub>50</sub>=10.41 ± 3.08 μM<br>k<sub>2</sub>/K<sub>I</sub> = 272 M<sup>-1</sup>s<sup>-1</sup> | IC<sub>50</sub>=9.0 ± 0.9 μM<br>K<sub>I</sub> = 2.29 μM |

<sup>a</sup>Data from reference.<sup>17</sup>

### Boceprevir, calpain inhibitors II and XII, and GC-376 have broad-spectrum antiviral activity against different CoVs

Given the binding and enzymatic inhibition of boceprevir, calpain inhibitors II and XII, and GC-376 against M^pro^ from multiple CoVs, we expect these compounds will have broad-spectrum antiviral activity against CoVs in cell culture. For this, cellular antiviral assays were performed against multiple human CoVs including HCoV-229E, HCoV-NL63, HCoV-OC43, MERS-CoV and SARS-CoV. Remdesivir was included as a positive control. It was found that all four compounds showed potent antiviral activity against all the CoVs tested in the viral cytopathic effect (CPE) assay in a dose-dependent manner (Figure 6). The 50% effective concentration EC_50_ values of GC-376 range from 99 nM to 3.37 µM (Figure 6 and Table 3). Calpain inhibitors II and XII have comparable potency as GC-376, with EC_50_ values in the range of 84 nM to 5.58 µM, and 100 nM to 1.97 µM, respectively (Figure 6 and Table 3). In contrast, boceprevir showed moderate antiviral activity against all the CoVs tested and the EC_50_ values were over 10 µM in most cases, except in the inhibition of SARS CoV-2 (EC_50_=1.31 ± 0.58 µM) (Figure 6 and Table 3). This result is in line with the weaker enzymatic inhibition of boceprevir against M^pro^s compared with GC-376, calpain inhibitors II and XII (Table 2).

**Figure 6.**
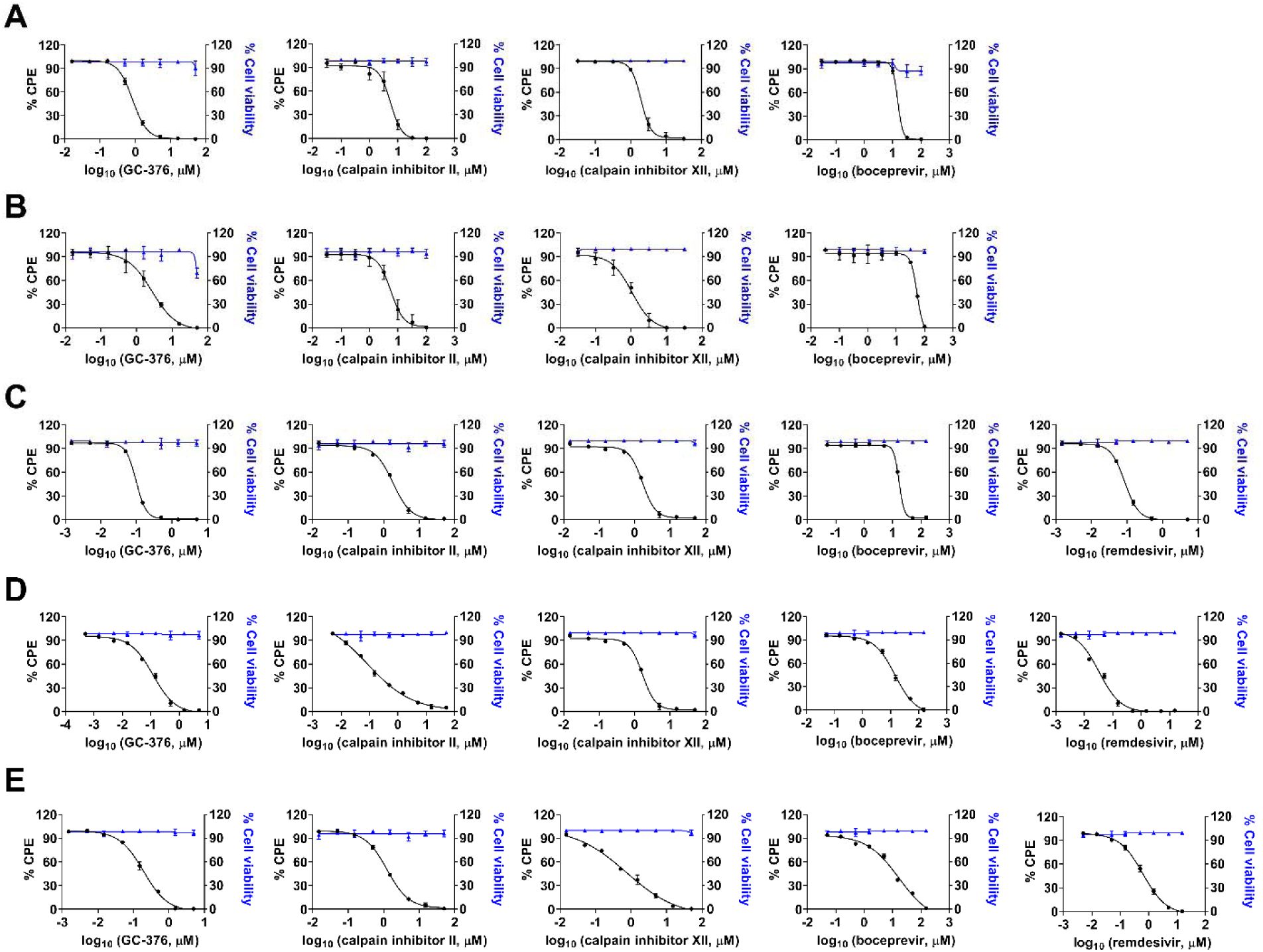
Cellular antiviral activity of boceprevir, calpain inhibitors II and XII, and GC-376 against MERS-CoV (A); SARS-CoV (B); HCoV-229E (C); HCoV-NL63 (D); and HCoV-OC43 (E) in CPE assay. Remdesivir was included in HCoV-229E, HCoV-NL63 and HCoV-OC43 CPE assays as a positive control. EC_50_ curve fittings for each compound in the CPE assay were obtained using log (concentration of inhibitors) vs percentage of CPE with variable slopes in prism 8. The cellular cytotoxicity test for each compound was included in each experiment and the resulting curves were shown in blue. Data are mean ± standard deviation of three replicates.

**Table 3.**
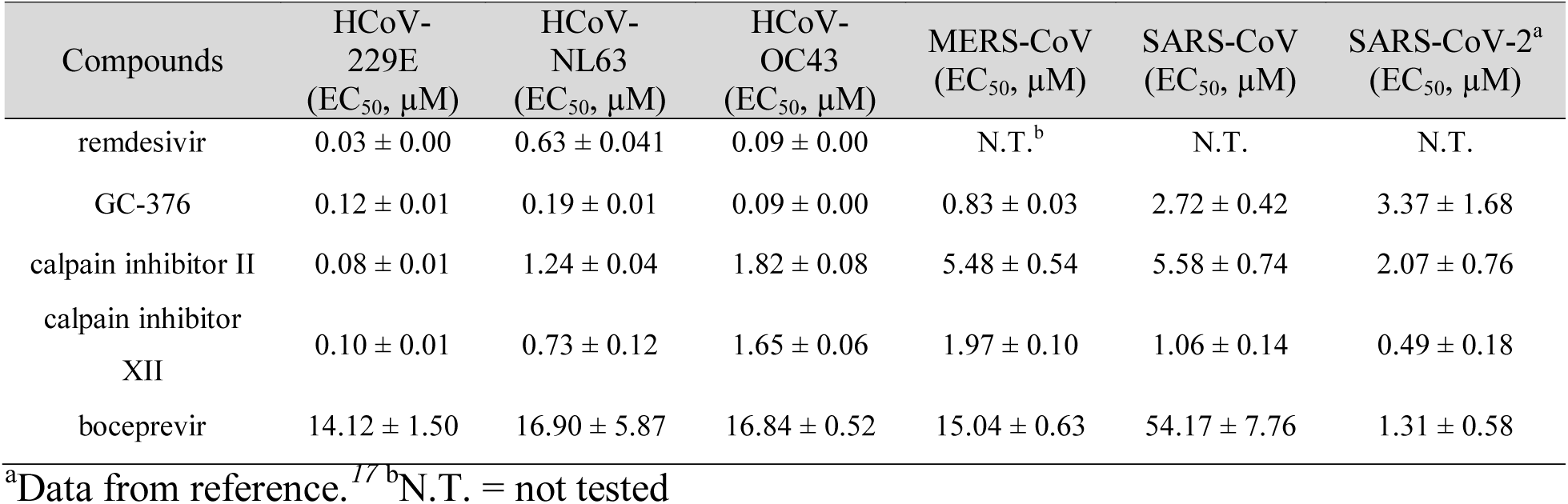
Broad-spectrum antiviral activity of boceprevir, calpain inhibitors II and XII, GC-376 against a panel of human CoVs in CPE assay.

| Compounds | HCoV-229E<br>(EC<sub>50</sub>, μM) | HCoV-NL63<br>(EC<sub>50</sub>, μM) | HCoV-OC43<br>(EC<sub>50</sub>, μM) | MERS-CoV<br>(EC<sub>50</sub>, μM) | SARS-CoV<br>(EC<sub>50</sub>, μM) | SARS-CoV-2<sup>a</sup><br>(EC<sub>50</sub>, μM) |
| --- | --- | --- | --- | --- | --- | --- |
| remdesivir | 0.03 ± 0.00 | 0.63 ± 0.041 | 0.09 ± 0.00 | N.T.<sup>b</sup> | N.T. | N.T. |
| GC-376 | 0.12 ± 0.01 | 0.19 ± 0.01 | 0.09 ± 0.00 | 0.83 ± 0.03 | 2.72 ± 0.42 | 3.37 ± 1.68 |
| calpain inhibitor II | 0.08 ± 0.01 | 1.24 ± 0.04 | 1.82 ± 0.08 | 5.48 ± 0.54 | 5.58 ± 0.74 | 2.07 ± 0.76 |
| calpain inhibitor XII | 0.10 ± 0.01 | 0.73 ± 0.12 | 1.65 ± 0.06 | 1.97 ± 0.10 | 1.06 ± 0.14 | 0.49 ± 0.18 |
| boceprevir | 14.12 ± 1.50 | 16.90 ± 5.87 | 16.84 ± 0.52 | 15.04 ± 0.63 | 54.17 ± 7.76 | 1.31 ± 0.58 |

<sup>a</sup>Data from reference.<sup>17</sup> <sup>b</sup>N.T. = not tested

To test whether boceprevir, calpain inhibitors II and XII, and GC-376 inhibit viral RNA synthesis, we performed the viral titer reduction assay using the RT-qPCR. HCoV-NL63 was chosen as a representative example and the viral nucleocapsid (N) gene expression level was detected in the absence or presence of different concentrations of compounds. Remdesivir was included as a positive control. All four compounds inhibited HCoV-NL63 viral N gene expression dose-dependently (Figure 7), giving EC_50_ values in the range of 0.96 to 19.86 µM (Figure 7), which were comparable to the EC_50_ values determined in the antiviral CPE assays (Table 3). Taken together, we have shown that boceprevir, calpain inhibitors II and XII, and GC-376 have broad-spectrum antiviral activity against CoVs.

**Figure 7.**
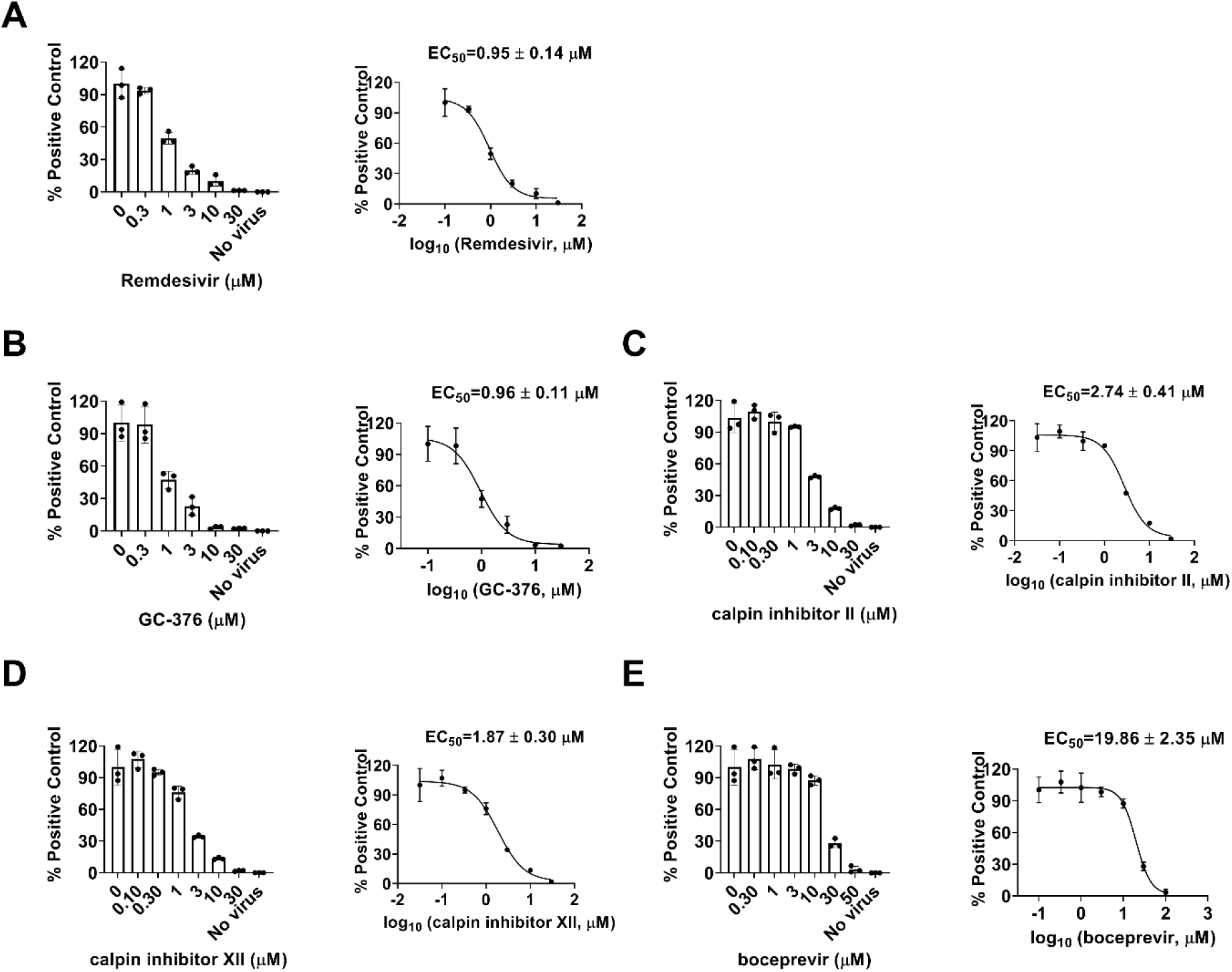
Dose-dependent inhibitory effect of boceprevir, calpain inhibitors II and XII, and GC-376 on HCoV-NL63 viral RNA synthesis in RT-qPCR assay. Positive control remdesivir (A); GC-376 (B); Calpain inhibitor II (C); Calpain inhibitor XII (D); and Boceprevir (E). The left figures represent the normalized RNA levels of the average of three repeats from each concentration tested, and EC_50_ curve fittings using log (concentration of inhibitors) vs normalized RNA levels with variable slopes in prism 8 were shown in the right figures. Data are mean ± standard deviation of three replicates.

### Combination therapy of boceprevir, calpain inhibitors II and XII, and GC-376 with remdesivir

Combination therapy has several advantages compared to monotherapies including delayed evolution of drug resistance, synergistic antiviral efficacy, and fewer side effects due to the lower amount of drugs used.^*28*^ Combination therapy has been extensively explored for the treatment in oncology, parasitic, bacterial and viral infections such as HIV and HCV.^*29–31*^ The combination treatment potentials of boceprevir, calpain inhibitors II and XII, and GC-376 with remdesivir were explored using HCoV-OC43 antiviral CPE assay. Combination indices (CIs) versus the EC_50_ values of compounds at different combination ratios were plotted as previously described.^*32*^ The red line indicates additive effect; the right upper area above the red line indicates antagonism, while the left bottom area below the line indicates synergy.^*32*^ In all combination scenarios, the combination indices at all the combination ratios fell on the red line (Figure 8), suggesting that boceprevir, calpain inhibitors II and XII, and GC-376 displayed an additive antiviral effect with remdesivir in the combination therapy.

**Figure 8.**
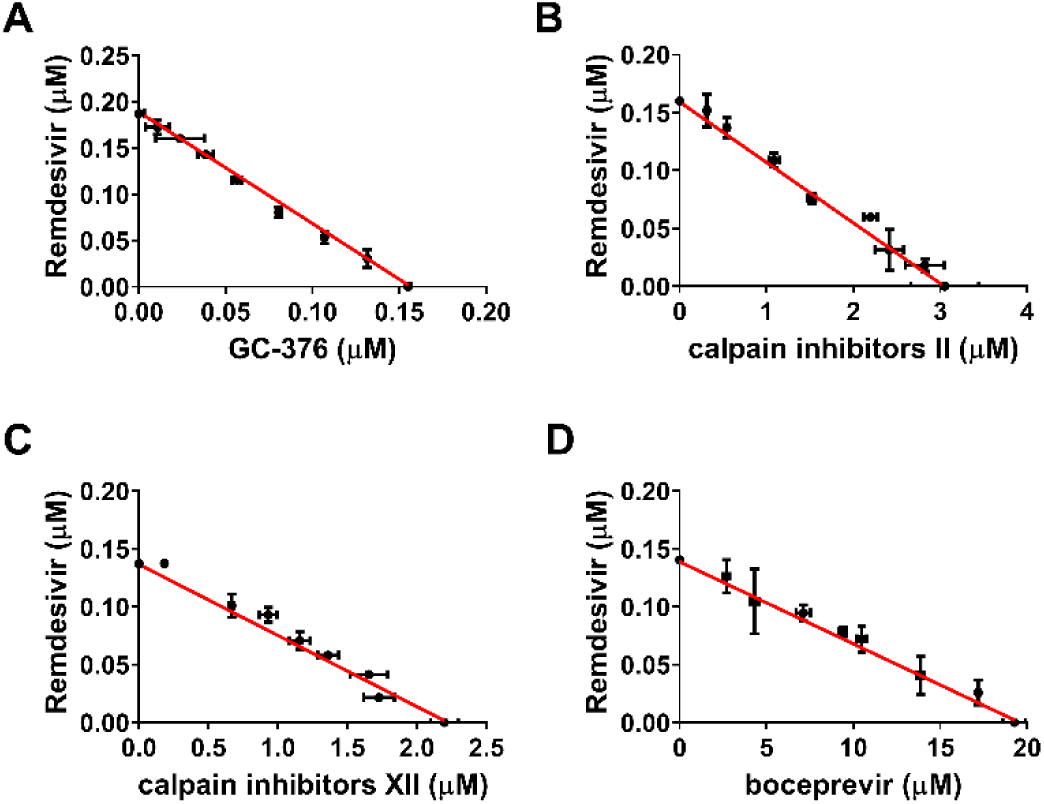
Combination therapy of remdesivir with GC-376 (A); Calpain inhibitor II (B); Calpain inhibitor XII (C); and Boceprevir (D). Data are mean ± standard deviation of three replicates.

## CONCLUSION

The COVID-19 pandemic emerged in late December 2019 in China has severe impacts on public health and global economy. It is one of the worst global pandemics in history. The high mortality rate and infection rate of COVID-19 is unprecedented. There has been three outbreaks of highly pathogenic and lethal human coronaviruses within the past two decades,^*3–5*^ and the current COVID-19 pandemic is a timely reminder of the urgent need for therapeutics of CoVs infection. In this work, we report the broad-spectrum antiviral activity of our previously identified SARS-CoV-2 M^pro^ inhibitors boceprevir, calpain inhibitors II and XII, and GC-376 against multiple CoVs including the highly pathogenic SARS-CoV, MERS-CoV, and human coronaviruses NL63, 229E, and OC43. Coupled with their antiviral activity against SARS-CoV-2 as reported earlier, this result suggests that these four compounds are promising drug candidates for the further development of broad-spectrum antivirals against not only current coronaviruses but also possibly future emerging coronaviruses. Among the four compounds, calpain inhibitors II and XII had a novel mechanism of action by targeting both the viral M^pro^ and host cathepsin L. In this study, we provided additional evidence from the pseudovirus neutralization assay as well as the drug time-of-addition assay to support the contribution of cathepsin L inhibition to the potent cellular antiviral activity of calpain inhibitors II and XII. We further demonstrated the additive antiviral effect of boceprevir, calpain inhibitors II and XII, and GC-376 with remdesivir in the combination therapy experiment.

It is known that CoVs enter host cells through two distinct pathways: the early membrane fusion pathway and the late endosomal pathway. For the early membrane fusion pathway, the transmembrane protease serine 2 (TMPRSS2) is responsible for the viral spike protein cleavage and activation.^*33*^ In the late endosomal entry pathway, the cysteine protease cathepsin L mediates the cleavage of spike protein.^*19*^ Several CoVs including SARS-CoV-2, SARS-CoV, MERS-CoV, HCoV-OC43, HCoV-229E, and HCoV-NL63 utilize cathepsin L for host cell entry,^*19, 24, 34–37*^ which offers an opportunity to develop broad-spectrum antivirals by targeting the cathepsin L protease. Indeed, cathepsin L inhibitors have been actively explored for the development of coronavirus antivirals.^*38–40*^ Compared with mono-specific M^pro^ inhibitors, the dual inhibitors such as calpain inhibitors II and XII that target both M^pro^ and cathepsin L might have additional advantages of synergistic antiviral effect and also possibly increased genetic barrier to drug resistance.

Although M^pro^ is relatively conserved among coronaviruses and picornaviruses, not all 3CL^pro^ or 3C^pro^ inhibitors have broad-spectrum antiviral activity. For example, the well-known 3CL^pro^ inhibitor rupintrivir inhibits the HCoV-229E with an EC_50_ value of 0.3 µM,^*41*^ however it failed to inhibit the M^pro^ from both SARS-CoV and SARS-CoV-2 M^pro^.^*17, 41*^ This might be due to the slight differences of residues located at the active sites of M^pro^. Gratifyingly, we have shown in this study that boceprevir, calpain inhibitors II and XII, and GC-376 inhibited multiple M^pro^s from different members of the coronavirus family and had potent cellular antiviral activity against all the coronaviruses tested. Among these four compounds, calpain inhibitors II and XII, and GC-376 had single-digit to submicromolar antiviral potency with a high selectivity index, while boceprevir had moderate antiviral activity. Nevertheless, boceprevir represents a novel chemotype that warrants the further development. Moving forward, continuous optimization of boceprevir, calpain inhibitors II and XII, and GC-376 might yield clinical candidates with favorable pharmacokinetic properties and proven *in vivo* antiviral efficacy in animal models. Such studies are ongoing and will be reported when available.

## METHODS

### Cell lines and viruses

The following reagent was obtained through BEI Resources, NIAID, NIH: Human Embryonic Kidney Cells (HEK-293T) Expressing Human Angiotensin-Converting Enzyme 2, HEK-293T-hACE2 Cell Line, NR-52511; SARS-Related Coronavirus 2, Wuhan-Hu-1 Spike-Pseudotyped Lentiviral Kit, NR-52948. Human rhabdomyosarcoma (RD), Vero E6, Huh-7, HEK293T expressing ACE2 (ACE2/293T), and HCT-8 cell lines were maintained in Dulbecco’s modified Eagle’s medium (DMEM); Caco-2 and MRC-5 cell lines were maintained in Eagle’s Minimum Essential Medium (EMEM). Both media were supplemented with 10% fetal bovine serum (FBS) and 1% penicillin-streptomycin antibiotics. Cells were kept at 37°C incubator in a 5% CO_2_ atmosphere. The following reagents were obtained through BEI Resources, NIAID, NIH: Human Coronavirus, OC43, NR-52725; Human Coronavirus, NL63, NR-470. HCoV-OC43 was propagated in RD cells; HCoV-NL63 was propagated in MRC-5 cells. HCoV-229E was obtained from Dr. Bart Tarbet (Utah State University) and amplified in Huh-7 or MRC-5 cells; The Urbani strain of severe acute respiratory syndrome coronavirus (SARS-CoV) and the EMC/2012 strain Middle East respiratory syndrome coronavirus (MERS-CoV) were obtained from the Centers for Disease Control and Prevention. Vero 76 cells were obtained from the American Type Culture Collection.

### Pseudovirus neutralization assay

A pseudotype HIV-1-derived lentiviral particles bearing SARS-CoV-2 spike and a lentiviral backbone plasmid encoding luciferase as reporter was produced in HEK293 T cells engineered to express the SARS-CoV-2 receptor, ACE2 (ACE2/293 T cells), as previously described.^*21*^ The pseudovirus was then used to infect Vero E6 cells in 96-well plates in the presence of DMSO or serial concentrations of E-64d, boceprevir, calpain inhibitors II and XII, and GC-376. At 48 hpi, cells from each well were lysed using the Bright-Glo Luciferase Assay System (Cat#: E2610, Promega, Madison, WI, USA), and the cell lysates were transferred to 96-well Costar flat-bottom luminometer plates. The relative luciferase units (RLUs) in each well were detected using Cytation 5 Cell Imaging Multi-Mode Reader (BioTek, Winooski, VT, USA).

### Drug time-of-addition experiment

A time-of-addition experiment was performed as previously described.^*42*^ Briefly, RD cells were seeded at 1 ×10^5^ cells/well in 12-well plate. 10 µM GC-376 or 50 µM calpain inhibitor XII was added at different time points, as illustrated in Figure 3. RD cells were infected with HCoV-OC43 at MOI of 0.1 at 24 h after seeding. Viruses in the supernatant were harvested at 16 hpi. The virus titers were quantified by plaque reduction assay.

### Differential scanning fluorimetry (DSF)

The binding of boceprevir, calpain inhibitors II and XII, and GC-376 on MERS-CoV, SARS-CoV and HCoV-OC43 M^pro^ proteins was monitored by differential scanning fluorimetry (DSF) using a Thermal Fisher QuantStudio™ 5 Real-Time PCR System as previously described^*43*^ with minor modifications. TSA plates were prepared by mixing M^pro^ proteins (final concentration of 4 μM) with different concentrations of compounds (0.2 to 200 µM) and incubated at 30 °C for 1 hr. 1× SYPRO orange (Thermal Fisher) were added and the fluorescence signal was recorded under a temperature gradient ranging from 20 to 95 °C (incremental steps of 0.05 °C/s). The melting temperature (*T*_*m*_) was calculated as the mid-log of the transition phase from the native to the denatured protein using a Boltzmann model in Protein Thermal Shift Software v1.3. Δ*T*_*m*_ was calculated by subtracting reference melting temperature of proteins in the presence of DMSO from the *T*_*m*_ in the presence of compounds. Curve fitting was performed using the Boltzmann sigmoidal equation in Prism (v8) software.

### Enzymatic assays

For the measurements of *K*_*M*_/*V*_*max*_ and IC_50_ values, proteolytic reactions were carried out with 100 nM MERS-CoV, SARS-CoV or HCoV-OC43 M^pro^ in 100 µL of pH 6.5 reaction buffer (20 mM HEPES, pH 6.5, 120 mM NaCl, 0.4 mM EDTA, 4 mM DTT and 20% glycerol) at 30 °C in a Cytation 5 imaging reader (Thermo Fisher Scientific) with filters for excitation at 360/40 nm and emission at 460/40 nm. Reactions were monitored every 90 s. For *K*_*M*_/*V*_*max*_ measurements, a FRET substrate concentration ranging from 0 to 200 µM was applied. The initial velocity of the proteolytic activity was calculated by linear regression for the first 15 min of the kinetic progress curves. The initial velocity was plotted against the FRET concentration with the classic Michaelis–Menten equation in Prism 8 software. For IC_50_ measurements, 100 nM M^pro^ protein was incubated with 0.1 to 100 µM boceprevir, calpain inhibitors II and XII, and GC-376 at 30 °C for 30 min in reaction buffer, then the reaction was initiated by adding 10 µM FRET substrate. The reaction was monitored for 1 h, and the initial velocity was calculated for the first 15 min by linear regression. The IC_50_ was calculated by plotting the initial velocity against various concentrations of the compounds using a dose-response curve in Prism 8 software. Kinetics measurements of the proteolytic reaction progress curves with boceprevir, calpain inhibitor II and XII, and GC-376 were carried out as follows: 60 nM MERS-CoV M^pro^, 5 nM SARS-CoV M^pro^, or 3.3 nM HCoV-OC43 M^pro^ was added to 20 µM FRET substrate with various concentrations of compounds in 200 µL of reaction buffer at 30 °C to initiate the proteolytic reaction. The reaction was monitored for 4 h. The progress curves were fitted as previously described.^*17*^ Substrate depletion was observed when proteolytic reactions progress longer than 90 min, therefore only the first 90 min of the progress curves were used in the curve fitting. In this study, k_2_ for boceprevir, calpain inhibitor II and XII, could not be accurately determined because of significant substrate depletion before the establishment of the equilibrium between EI and EI*, leading to very slow k_2_. In these cases, K_I_ was determined with Morrison equation in Prism 8.

### SARS-CoV and MERS-CoV CPE assays

Antiviral activities of test compounds were determined in nearly confluent cultures of Vero 76 cells. The assays were performed in 96-well Corning microplates. Cells were infected with approximately 30 cell culture infectious doses (CCID_50_) of SARS-CoV or 40 CCID_50_ of MERS-CoV. The plates were incubated at 37°C with 5% CO_2_ and 50% effective concentrations (EC_50_) were calculated based on virus-induced cytopathic effects (CPE) quantified by neutral red dye uptake after 4 days of incubation for SARS-CoV or 3 days of incubation for MERS-CoV. Three microwells at each concentration of compound were infected. Two uninfected microwells served as toxicity controls. Cells were stained for viability for 2 h with neutral red (0.11% final concentration). Excess dye was rinsed from the cells with phosphate-buffered saline (PBS). The absorbed dye was eluted from the cells with 0.1 ml of 50% Sörensen’s citrate buffer (pH 4.2)-50% ethanol. Plates were read for optical density determination at 540 nm. Readings were converted to the percentage of the results for the uninfected control using an Excel spreadsheet developed for this purpose. EC_50_ values were determined by plotting percent CPE versus log10 inhibitor concentration. Toxicity at each concentration was determined in uninfected wells in the same microplates by measuring dye uptake.

### HCoV-OC43, HCoV-229E, and HCoV-NL63 CPE assays

Antiviral activities of boceprevir, calpain inhibitors II and XII, and GC-376 against HCoV-229E, HCoV-NL63 and HCoV-OC43 were tested in CPE assays as previously described^*44*^ with minor modifications. Briefly, cell cultures near confluency in 96-well plates were infected with 100 µL of viruses at desired dilutions and incubated for 1 h. Unabsorbed virus was removed and different concentrations of testing compounds (0, 0.01, 0.03, 0.1, 0.3, 1, 3, 10, 30, 100 µM) were added. Remdesivir was included as a positive control. The plates were incubated for another 3 to 5 days when a significant cytopathic effect was observed in the wells without compound (virus only). Cells were stained with 0.1 mg/ml neutral red for 2 h, and excess dye was rinsed from the cells with PBS. The absorbed dye was dissolved with a buffer containing 50% ethanol and 1% glacial acetic acid. Plates were read for optical density determination at 540 nm. Readings were normalized with uninfected controls. EC_50_ values were determined by plotting percent CPE versus log_10_ compound concentrations from best-fit dose response curves with variable slope in Prism 8. Toxicity at each concentration was determined in uninfected cells in the same microplates by measuring neutral red dye uptake.

### RNA extraction and real-time PCR

RNA extraction and RT-PCR were performed as previously described.^*45*^ Total RNA was extracted from HCoV-NL63 virus infected Caco-2 cells at a MOI of 0.05 at 48 hours post infection using TRIzol reagents (Thermo Fisher Scientific). 2.0□μg of total RNA was used to synthesize the first strand cDNA of viral RNA and host RNA using SuperScript III reverse transcriptase (Thermo Fisher Scientific) and Random Hexamer primer. Viral RNA was amplified on a Thermal Fisher QuantStudio™ 5 Real-Time PCR System (Thermo Fisher Scientific) using FastStart Universal SYBR Green Master mix (carboxy-X-rhodamine; Roche) and HCoV-NL63 N gene-specific primers (Forward: 5’-CTGTTACTTTGGCTTTAAAGAACTTAGG-3’; Reverse: 5’-CTCACTATCAAAGAATAACGCAGCCTG-3’). GAPDH was also amplified to serve as a control using human GAPDH-specific primers (GAPDH-F: 5′-ACACCCACTCCTCCACCTTTG-3′ and GAPDH-R: 5′-CACCACCCT GTTGCTGTAGCC-3′). The amplification conditions were: 95□°C for 10□min; 40 cycles of 15□s at 95□°C and 60□s at 60□°C. Melting curve analysis was performed to verify the specificity of each amplification. All experiments were repeated three times independently.

### Combination therapy

Boceprevir, calpain inhibitors II and XII, and GC-376 was mixed with remdesivir at combination ratio of 8:1, 4:1, 2:1, 1:1, 1:2, 1:4, and 1:8 separately. The mixture of each compound with remdesivir at each combination ratio was serially diluted into 7 different concentrations and applied in HCoV-OC43 CPE assay to determine EC_50_ of each compound and remdesivir in the combination ratio. A combination indices (CIs) plot was used to depict the EC_50_ values of each compound and remdesivir at different combination ratios. The red line indicates the additive effect, and above the red line indicates the antagonism, while below the red line indicates the synergy.^*32*^

## Author Information

### Author Contributions

Y.H., C.M., and J.W. designed the experiments. Y.H. performed the enzymatic assay, thermal shift binding assay, time-of-addition assay, pseudovirus neutralization assay, antiviral assays against HCoV-OC43, HCoV-229E, and HCoV-NL63, and the combination therapy experiment. C.M. performed the enzymatic assays against SARS-CoV-2 M^pro^ and MERS-CoV M^pro^. T. S. expressed and purified the SARS-CoV-2, SARS-CoV, and MERS-CoV M^pro^. B. H. and B. T. performed the SARS-CoV and MERS-CoV CPE assays. Y.H. and J.W. wrote the manuscript.

## Acknowledgments

This research is supported by the NIH grants (AI119187, AI144887, AI147325, and AI157046), and the Young Investigator Award grant from the Arizona Biomedical Research Centre to J.W (ADHS18-198859).

## Table of Content Graphic

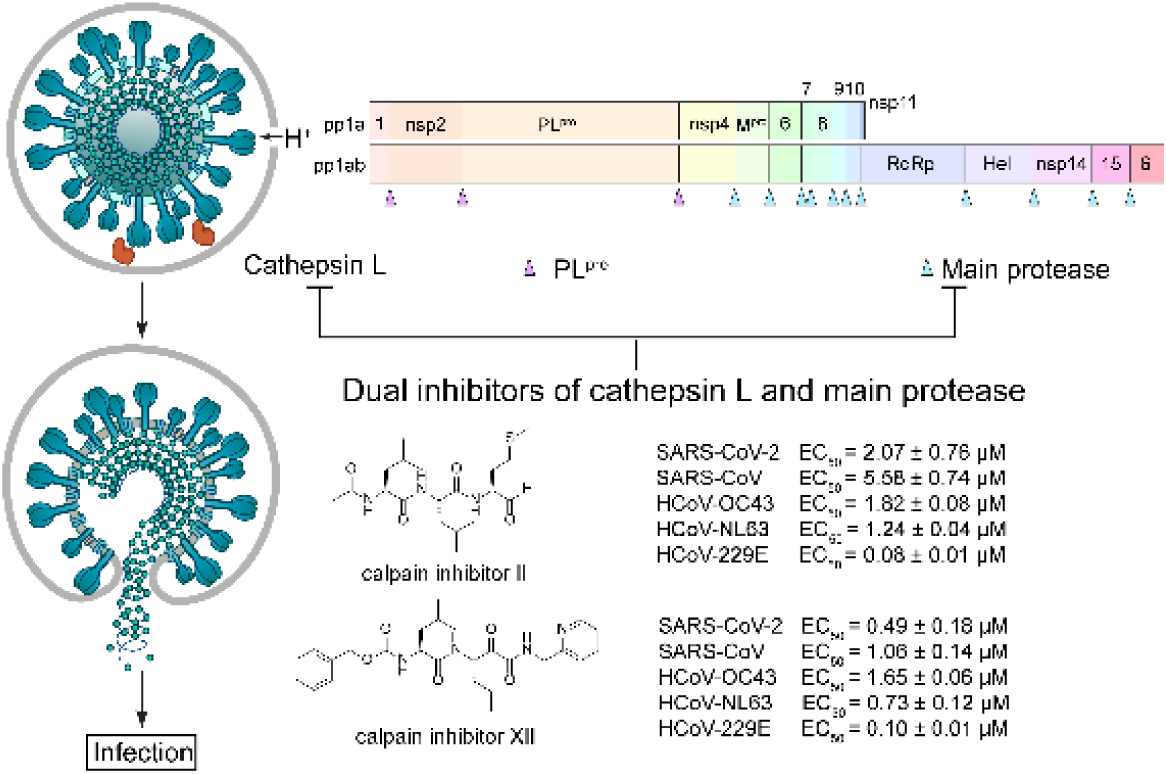

