## Supplementary Information for "Boceprevir, calpain inhibitors II and XII, and GC-376 have broad-spectrum antiviral activity against coronaviruses in cell culture"

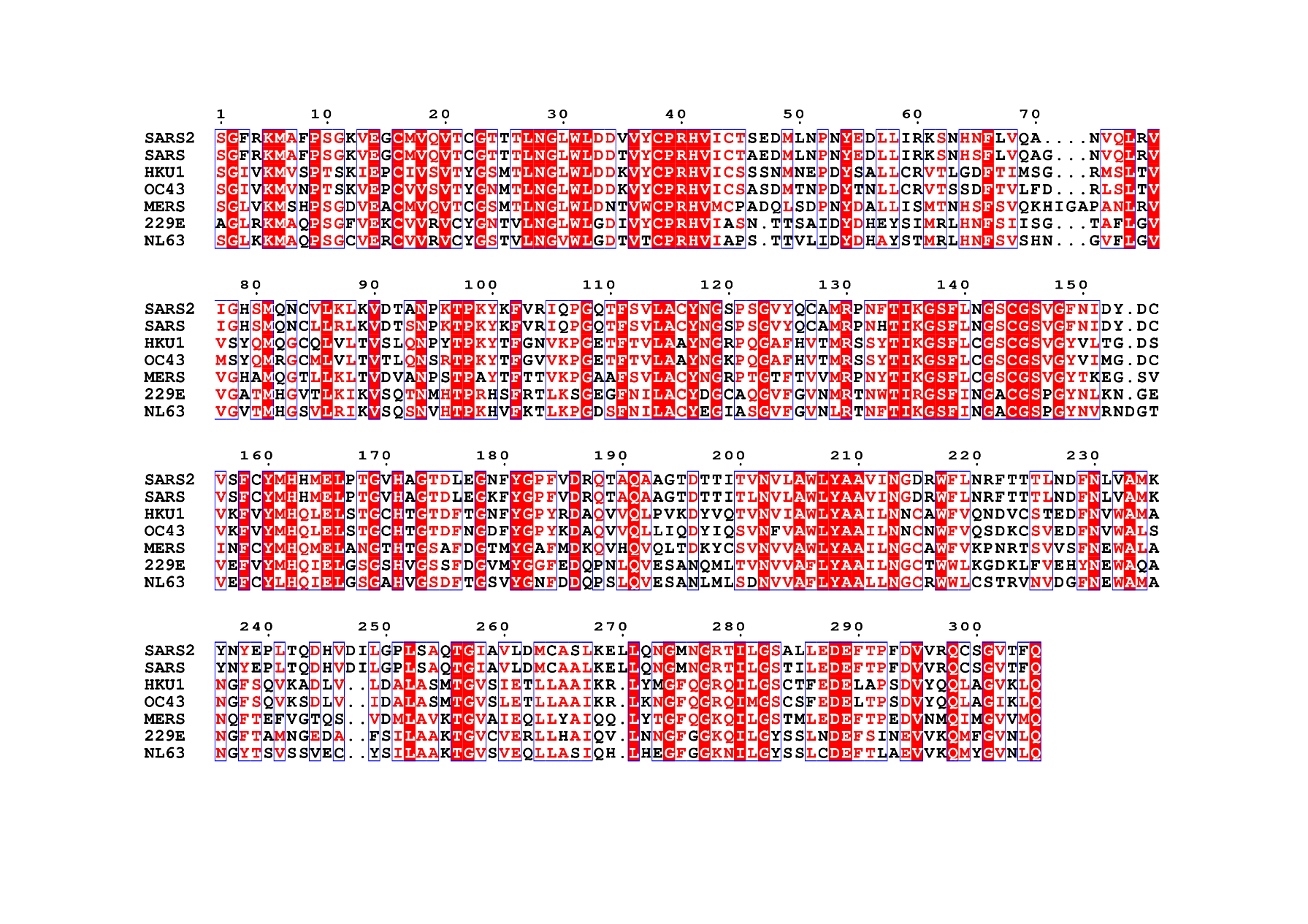

**Figure S1.** Amino acid sequence alignment of M^pro^ from SARS-CoV-2 (Accession No.: 6WQF_A), SARS-CoV (Accession No.: 2C3S_A) MERS-CoV (Accession: 5C3N_A) HCoV-229E (Accession No.: 2ZU2_B) HCoV-NL63 (Accession No.: 3TLO_B) HCoV-OC43 (Accession No.: YP_009924323) HCoV-HKU1 (Accession No.: YP_459936). The protein sequences were downloaded from the National Center for Biotechnology Information (NCBI).

Table S1. Amino acid conservation between the seven HCoVs shown alignment in Figure S1.

|  | | SARS-CoV-2 | SARS-CoV | MERS-CoV | HCoV-HKU1 | HCoV-229E | HCoV-OC43 | HCoV-NL63 |
| --- | --- | --- | --- | --- | --- | --- | --- | --- |
| SARS-CoV-2 | Identity% |  | 95 | 49 | 48 | 40 | 48 | 43 |
|  | Similarity% |  | 98 | 67 | 64 | 60 | 66 | 63 |
| SARS-CoV | Identity% | 95 |  | 50 | 48 | 39 | 48 | 42 |
|  | Similarity% | 98 |  | 67 | 64 | 60 | 66 | 62 |
| MERS-CoV | Identity% | 49 | 50 |  | 54 | 48 | 52 | 47 |
|  | Similarity% | 67 | 67 |  | 68 | 65 | 69 | 65 |
| HCoV-HKU1 | Identity% | 48 | 48 | 54 |  | 45 | 82 | 43 |
|  | Similarity% | 64 | 64 | 68 |  | 64 | 92 | 63 |
| HCoV-229E | Identity% | 40 | 39 | 48 | 45 |  | 44 | 70 |
|  | Similarity% | 60 | 60 | 65 | 64 |  | 64 | 84 |
| HCoV-OC43 | Identity% | 48 | 48 | 52 | 82 | 44 |  | 42 |
|  | Similarity% | 66 | 66 | 69 | 92 | 64 |  | 62 |
| HCoV-NL63 | Identity% | 43 | 42 | 47 | 43 | 70 | 42 |  |
|  | Similarity% | 63 | 62 | 65 | 63 | 84 | 62 |  |

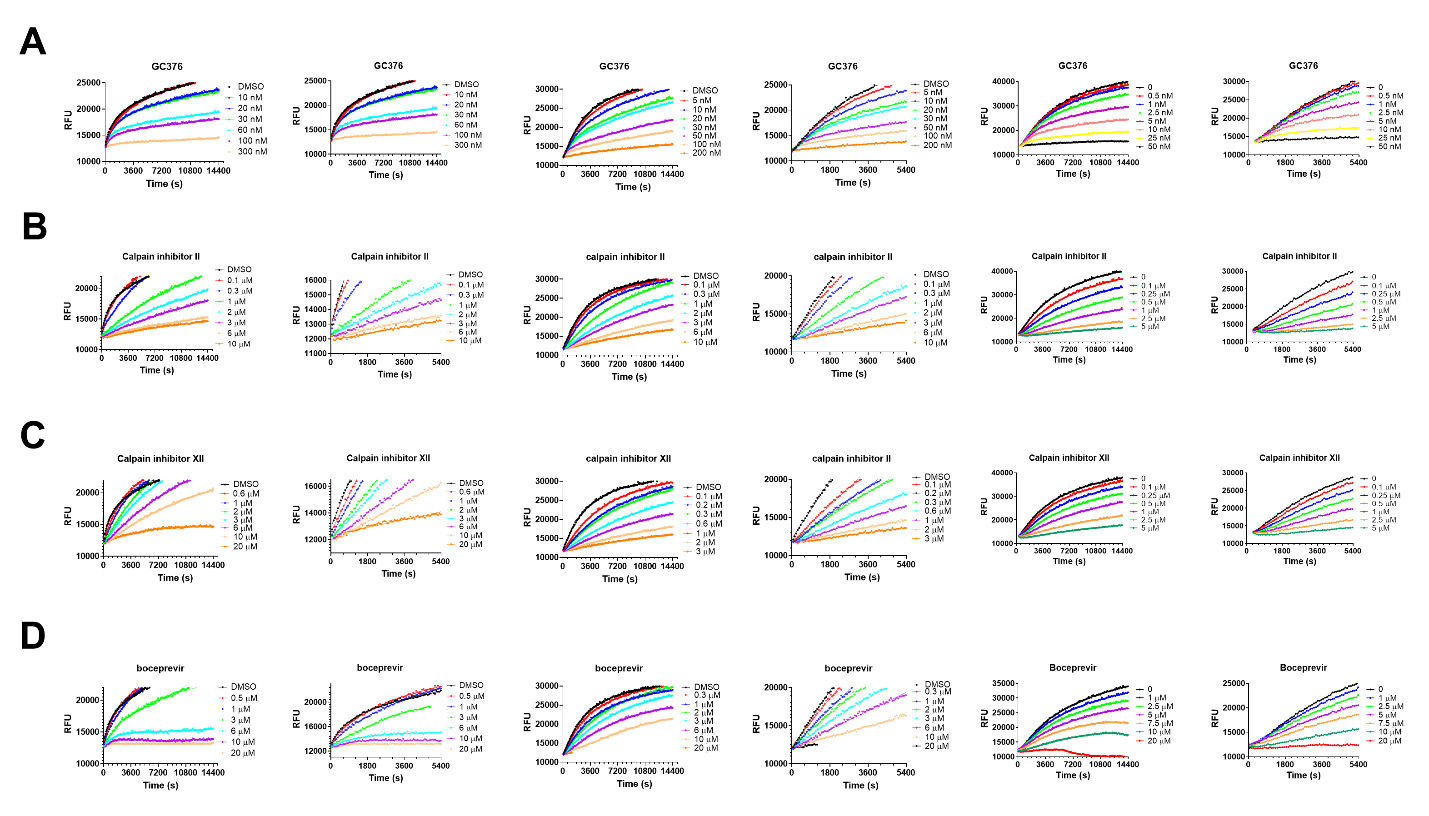

**Figure S2.** Proteolytic reaction progression curves of MERS-CoV M^pro^ (left two columns), SARS-CoV M^pro^ (middle two columns) and HCoV-OC43 M^pro^ (right two columns) in the presence or the absence of compounds. In the kinetic studies, 60 nM MERS-CoV M^pro^, 5 nM SARS-CoV M^pro^ or 3.3 nM HCoV-OC43 M^pro^ was added to a solution containing various concentrations of protease inhibitors and 20 µM FRET substrate to initiate the reaction. The reaction was then monitored for 4 h. For each M^pro^ tested, the left column shows the reaction progression up to 4 h; right column shows the progression curves for the first 90 min, which were used for curve fitting to generate the plot shown in Figure 4. Detailed methods were described in “Materials and methods” section. GC-376 (A); calpain inhibitor II (B); calpain inhibitor XII (C); boceprevir (D).
